# Pillar arrays as tunable interfacial barriers for microphysiological systems

**DOI:** 10.1101/2025.01.08.632020

**Authors:** Ishan Goswami, Yongdeok Kim, Gabriel Neiman, Brian Siemons, Jazmin I. Velazquez, Kerem Yazgan, Tammy Ng, Kevin E. Healy

**Affiliations:** Department of Bioengineering, University of California Berkeley; Department of Materials Science and Engineering, University of California Berkeley

## Abstract

We report on the design and fabrication of a novel circular pillar array as an interfacial barrier for microfluidic microphysiological systems (**MPS**). Traditional barrier interfaces, such as porous membranes and microchannel arrays, present limitations due to inconsistent pore size, complex fabrication and device assembly, and lack of tunability using a scalable design. Our pillar array overcomes these limitations by providing precise control over pore size, porosity, and hydraulic resistance through simple modifications of pillar dimensions. Serving as an interface between microfluidic compartments, it facilitates cell aggregation for tissue formation and acts as a tunable diffusion barrier that mimics diffusion in vivo. We demonstrate the utility of barrier design to engineer physiologically relevant cardiac microtissues and a heterotypic model with vasculature within the device. Its tunable properties offer significant potential for drug screening/testing and disease modeling, enabling comparisons of drug permeability and cell migration in MPS tissue with or without vasculature.

---

Microfluidics has been widely adopted to create microphysiological systems (**MPS**) that can be used to culture cells/tissues in near-physiological conditions via the provision of dynamic flow conditions, physiological ratios of tissue and culture medium, and the ability to create 3D tissue geometries. We and others have shown the application of these devices for drug testing and disease modeling.^1–8^ Many of these microfluidic devices consist of a tissue chamber separated from the media channels via a diffusion barrier that controls the transport rate of biomolecules to and from the tissues. Two widely used form factors of engineered interfaces as diffusion barriers are porous membranes^9–11^ and microchannel barriers.^12–16^

Thin porous polymeric membranes have been used to separate the tissue chamber from the media channels.^9–11^ The porous membrane allows the diffusion of nutrients and molecules as well as allowing heterotypic cell-cell communication via the incorporation of two different cell-types on either side of the membrane. The advantage of this approach is the commercial availability of porous membranes made of polymers such as polyethylene terephthalate and polycarbonate. The porosity in these polymeric membranes is achieved via a track-etching process involving bombardment of polymer with heavy ions followed by chemical etching.^17^ The fabrication process is scalable and allows for wide adoption. However, there is huge variation in pore sizes and pore position in these membranes. Furthermore, the process of incorporating these membranes into a MPS involves functionalizing membranes and cumbersome alignment with the microfluidic chambers during assembly.^1,2^

Another form-factor uses an array of microchannels as a diffusion barrier separating the media channels and the tissue chamber.^12–16^ The hydraulic diameter of the microchannels provides the equivalent pore-size achieved via polymeric membranes. These microchannel-based barriers are incorporated into the design of the master-mold used to create the polymeric MPS device and therefore minimizes any cumbersome alignment and additional functionalization during assembly. Furthermore, the pore-size and position of the microchannels are easily controlled during the design and fabrication of the master-mold. However, patterning of these microchannel arrays, including pitch and length of the channels, are subject to the aspect ratio and design of the tissue chamber. This can sometimes be limiting while recreating heterotypic cell models across the barrier due to the discrete nature of the pores. Furthermore, a common procedure to load cells into the microfluidic tissue chambers involves the centrifugation of cells into the chamber or application of a vacuum, which results in filling of the tissue chambers. The diffusion barrier plays an important role in the successful loading of the chamber during the loading as they act as a burst-valve due to the air-water interface, thus providing a resistance during loading without cells escaping into the media channel. Thus, the geometric features of such microchannel arrays must accommodate a compromise between providing a hydraulic resistance for a successful filling of the tissue chamber and a diffusion barrier that allows biomolecular diffusion and ability to create heterotypic models across the barrier.

In this article, we demonstrate the development of a pillar array barrier in a MPS device that combines the advantages offered by porous membrane and microchannel barriers. We show that this pillar array diffusion barrier allows precise control over the pore location and the porosity that can be accurately tuned. The barrier does not require cumbersome handling and assembly. Furthermore, the polymeric pillar template allows control of the burst pressure and diffusion by simply changing the height of the array during fabrication. We hypothesized that control of burst pressure and diffusion by the pillar barrier allows robust tissue formation and provides an avenue to create physiologically dense tissue that models in vivo tissue physiology.

We implemented this barrier design in a cardiac MPS. We first present the concept of the barrier design, the fabrication procedure, and provide estimates of the burst pressure of different configurations of barriers. The ability to control biomolecular diffusion via alteration of the geometric parameters of the barrier is demonstrated via computational predictions made via the finite element method (**FEM**). Next, the ability of the barrier to accommodate the aggregation and formation of physiologically dense microtissue is demonstrated. The barrier allowed the exchange of nutrients to the microtissues at physiologically relevant dynamic flow. Lastly, we show how the pillar array can be used to form a barrier model that incorporates endothelial cells to “vascularize” a cardiac tissue.

## RESULTS AND DISCUSSION

### Design and fabrication of the barrier

The MPS barrier consists of an array of pillars (**Figure 1A**) separating two compartments, i.e. the cell chamber and the media channel. The porosity of the barrier can be altered by changing the distance between two pillars, defined in **Figure 1A** as pore size. In our design, the porosity is defined by the ratio of the volume of voids to the total volume. For example, for a simple rectangular space of 125 µm x 708 µm (see, **Figure S1A**), altering the pore size from 8 µm to 2 µm changes the porosity from 31% to 19% in a single configuration where the number of pillars is kept same (i.e. 8) and their pillar diameters altered to achieve the pore size. It is noted that porosity can be altered in multiple ways by varying the number of pillars and pore size determined via the pillar diameters. On the other hand, changing the pillar height allows altering the porous volume that controls the hydraulic resistance offered by the barrier to a biomolecule and/or cell. For example, keeping the pore size constant at 8 µm, the porous volume can be linearly varied by increasing or decreasing the pillar height. Thus, the pillar array barrier allows flexibility in tuning parameters such as porosity and hydraulic resistance that serve as a tunable barrier in an MPS.

**Figure 1:**
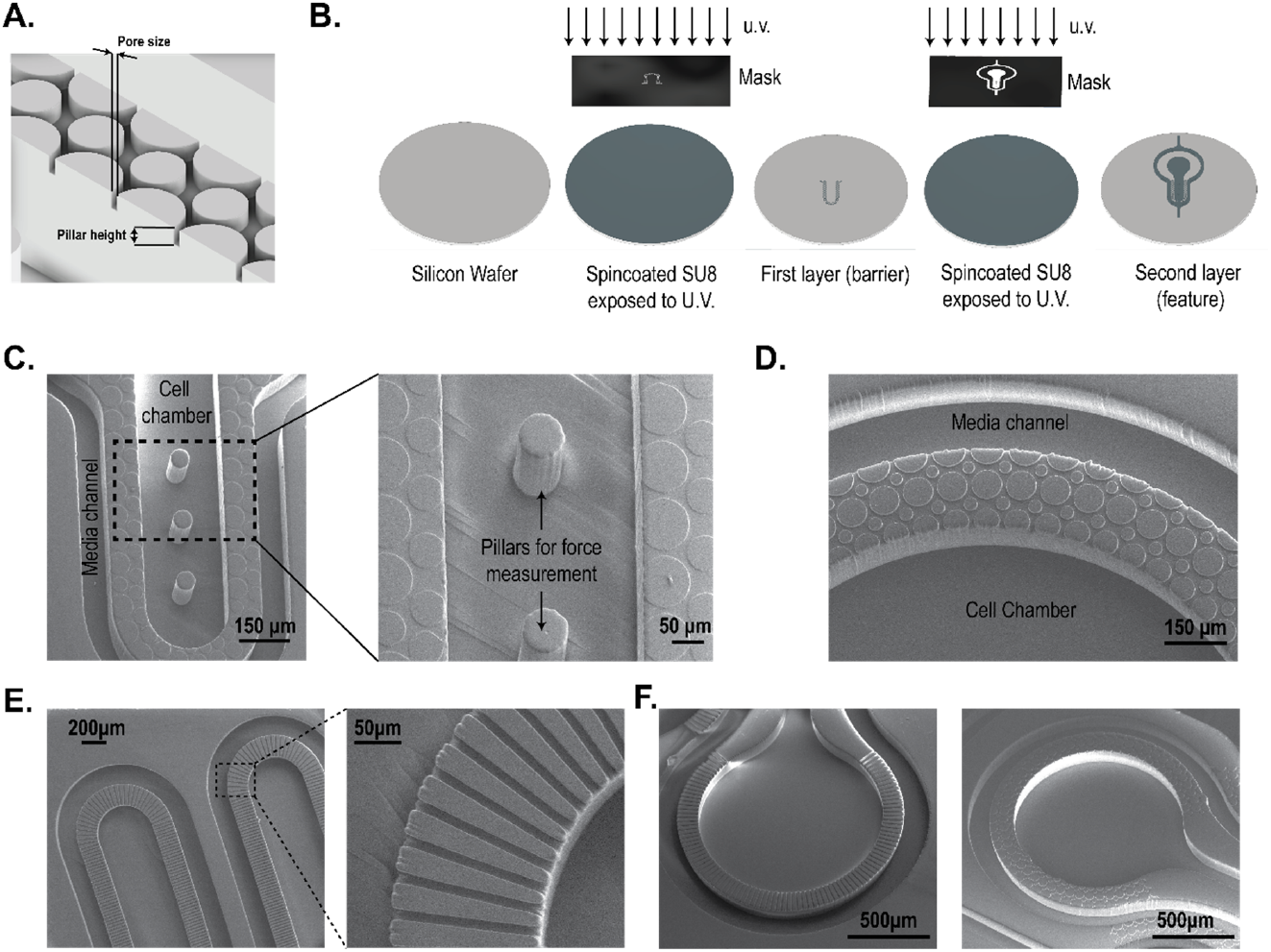
Fabrication of pillar array barrier MPS device. **(A)** An illustration of the replica molded pillar array barrier generated from the master mold. Pore size was defined in the mask design and the thickness of the first layer of spin-coated photoresist determined pillar height. **(B)** Illustrated process to create a multi-layered master mold using SU8 photochemistry. The photoresist was spin-coated on a silicon wafer and the first layer was generated by exposing it to u.v. light. After developing the first layer, fiduciary markers were used to create a second layer following a similar protocol of spin-coating and u.v. exposure. **(C)** SEM images of the PDMS replica molds of the cardiac MPS. A barrier separates the cell chamber and the media channel. Force-measuring pillars were fabricated within the cell chamber. Shown here is a barrier height of 2 µm, and cell chamber depth of 150 µm. **(D)** SEM image of replica mold with pillar array-based barriers of 10 µm pore size and 10 µm barrier height. The cell chamber and media channel are 100 µm in depth. **(E)** Example of a microchannel interface barrier in an MPS. **(F)** Comparison of a microchannel and a pillar array MPS interface barrier.

We demonstrate two designs using different pore sizes and pillar heights in an MPS device. As an example, **Figure 1C** shows the SEM of a cardiac MPS that follows a design form factor previously reported by our group.^3,12,18^ The major components of the device are a central cell chamber, two adjacent media channels, and the pillar array barrier. The cell chamber has a width of 300 µm, while the media channels are 100 µm. The height of both the cell chamber and media channel is approximately 150 µm. The cell chamber is connected to the media channels by the pillar array barrier of height 2 µm and pore size of 8 µm. Details of the fabrication protocol are provided in the Methodology section. **Figure 1D** shows another form factor where the pillar height and pore size were 10 µm, and separate a media channel (100 µm width) and a circular cell chamber of 1500 µm diameter. In both designs, the pillar array acts as a barrier to aggregate cells within the cell chamber while loading and as a diffusion barrier that can model the endothelial compartment separating the microtissue from the perfusion medium, protecting the tissue itself from shear forces but allowing nutrient exchange. As a comparison to these pillar array interfaces, **Figures 1E-F** show microchannel-based fenestrations.^12,15^

In the next subsections, we report the characterization of the pillar array barrier in the cardiac MPS (**Figure 1C**) via burst-pressure and biomolecular diffusion measurements.

### Pressure barrier offered by pillar array diffusion barrier

The loading of cells into the cardiac MPS cell chamber involves centrifugation of singularized cells or spheroids suspended in a cell culture medium. For a successful aggregation of the cells/spheroids into a microtissue, the barrier must act as a burst valve to allow the cells to be contained within the cell chamber. During the cell loading process, the cell medium replaces the air within the cell chamber and faces a high-pressure barrier at the opening of the pillar array due to an abrupt change in the cross-section area, stopping the liquid advancement. Thus, the diffusion barrier in the MPS acts as a capillary burst valve regulating the fluid flow. Empirically, the pressure required for the liquid to burst across a capillary burst valve (Δp_b_) depends on the liquid–air surface tension (*γ*_*lv*_) and the hydraulic diameter of the channel (D_h_), such that *Δp*_*b*_ *∝ (γ*_*lv*_/*D*_*h*_).^19–21^ The expression of burst pressure depends on the capillary valve geometry as well. This forms the basis the pillar array barrier, whereby variations in the circular geometry within a single fenestra achieve a higher burst pressure (**Figure 2A**) than a single microchannel-based straight fenestra (**Figure S1B**).

**Figure 2:**
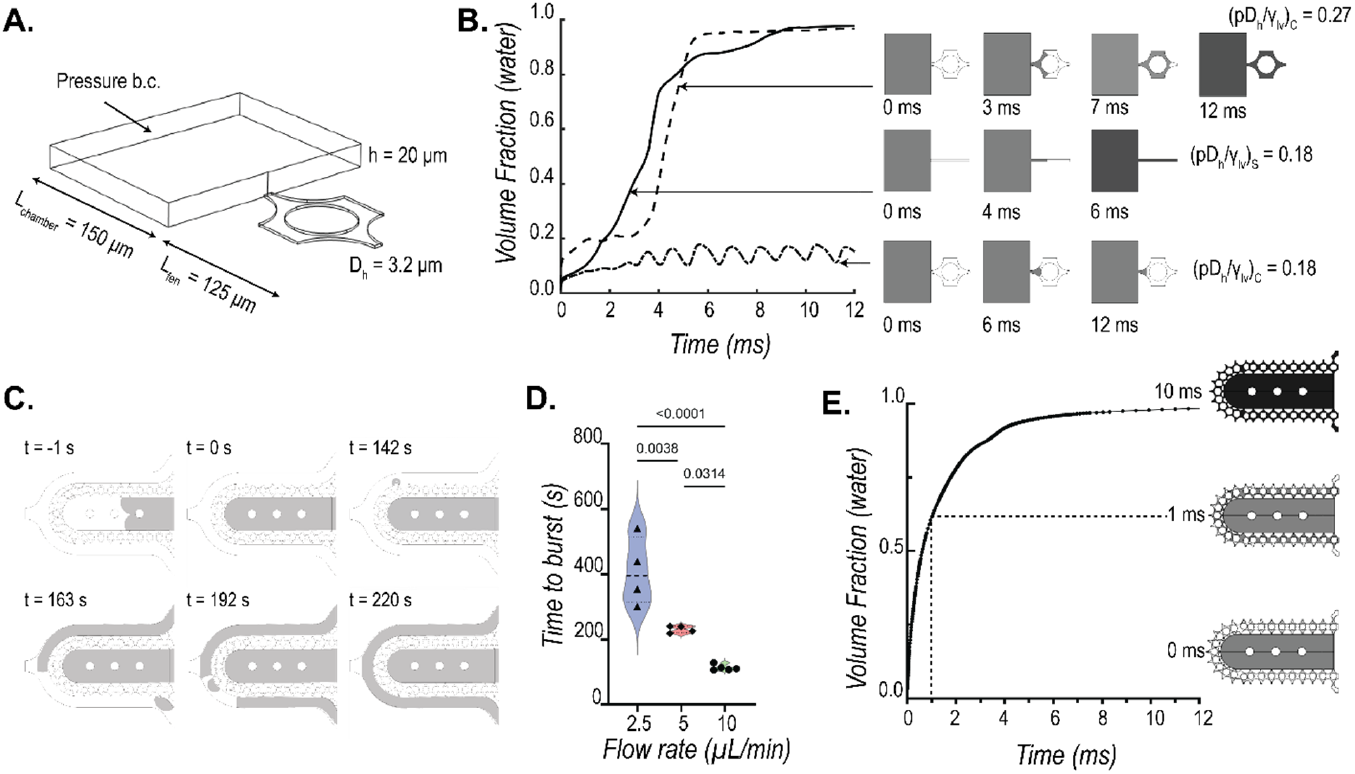
Characterization of burst pressure offered by the pillar array barrier. **(A)** A single fenestra finite element model was used to simulate the movement of the air-water interface across the microfluidic barrier to quantify burst pressure. **(B)** Numerically estimated transient water volume fraction in the single fenestra models upon application of pressure (grey: water; white: air). The comparison shows the pillar barrier has a higher burst pressure than a single straight microchannel barrier for the same hydraulic diameter (D_h_ = 3.2 µm). Interfacial properties *γ*_*lv*_ *=* 0.072 *N*/*m* and contact angle *θ*_*c*_ *=* 66° were used for the calculations. **(C)** Representative sequences of experimentally captured movement of air-water interface (grey: FITC-doped water; white: air) as water is pumped into the cell chamber in the MPS device. The time to first burst (shown at t= 142s) is used to quantify the barrier function **(D)** offered at three different flow rates. **(E)** Finite element model predicted that the bursting takes place within fractions of a second when the pressure within the cell chamber reaches a critical value. The trace of water volume fraction shown here was obtained when applied pressure at the cell chamber inlet was 6000 Pa. A fenestration height of 4 µm and a cell chamber height of 150 µm were used in this simulation.

To model the difference in resistance offered by a straight microchannel and a pillar array barrier, we implemented a level set method available via the COMSOL FEM package to simulate the air and water multiphase transport and calculate the burst pressures across different barrier designs. **Figure 2A** shows the computational domain used to simulate the air-water interface movement as we apply a constant pressure boundary condition to force water from a chamber (length: 150 µm, width: 200 µm, height: 20 µm) across a 125 µm length single fenestra of height 2 µm. The hydraulic diameters for straight microchannel and pillar array fenestra were kept at 3.2 µm. For numerical stability and convergence, the values for the interfacial properties were kept constant for both the geometries at *γ*_*lv*_=0.072 N/m and contact angle *θ*_*c*_= 66°. **Figure 2B** shows the transient volume fraction occupied by water in the fenestra as predicted by the level-set method for different applied pressures, normalized by *γ*_*lv*_/*D*_*h*_, to obtain a non-dimensional pressure *p\**. As seen from **Figure 2B**, a *p\** of 0.18 is enough to burst water out of the straight microchannel-based fenestra (*(pD*_*h*_/*γ*_*lv*_)_*s*_) but not sufficient to burst the valve formed by the pillar array. The pressure required to burst the pillar array was 1.5 times that required for the microchannel fenestra as seen from the *p\** of 0.27. The reason for this increased burst pressure can be explained by the pillar array fenestra consisting of a converging-diverging cross-sectional area that provides additional resistance for the moving liquid interface.

To validate the burst valve functionality of the pillar array experimentally, we recorded the movement of air-water interface as FITC-doped water was pumped into an empty cell chamber of the MPS. A representative sequence of these recordings is shown in **Figure 2C**. We pumped water at flow rates of 2.5 µL/min, 5 µL/min, and 10 µL/min. Time to burst was defined as the first signal of water in the media channel. Based on this criterion, we see a clear barrier function offered by the pillar arrays as shown by the differential time to bursts at the three flow rates (**Figure 2D**). It is noted that as opposed to computational study, our boundary condition here is a mass inflow condition rather than a pressure boundary condition. This explains the different timescales estimated to burst. The gradual increase of liquid mass within the cell chamber with the flow rates allows the build-up of pressure and once a critical pressure is reached, the liquid bursts across the barrier. This takes a few hundred seconds with a flow inlet condition. A fluctuation of the liquid advancing front in the capillary valve as recorded during our experiments is provided as reference in the supplementary materials (**SV1**). We see that when a high pressure of 6000 Pa is applied in our 3D level set model, the water interface bursts across the barrier without any fluctuations (**Figure 2E**). Overall, we can infer that pressure builds up with the gradual influx of liquid in the cell chamber. The moving liquid front within the barrier fluctuates until critical pressure is reached, at which point the liquid floods spontaneously.

### Characterization of molecular transport across the diffusion barrier

Since altering the pore size and pillar height of the barrier changes the porous volume, it allows us to tune the diffusion across the barrier. To characterize the molecular transport across the barrier, we simulated the transport of 70 kDa dextran when injected into the media channel at concentration of c_0_ (7 mmol/m^3^) and a rate of 80 µL/min. The temporal evolution of dextran was obtained by coupling the Navier-Stokes solver in COMSOL to solve for the flow field and the Transport of Diluted Species solver to determine the concentration profiles. **Figure 3A** shows the transient evolution of space-averaged value of dextran concentration at the middle of the cell chamber. We used 70 kDa dextran due to its large size and hence lower diffusivity (D= 45.8 µm^2^/s).^22,23^

**Figure 3:**
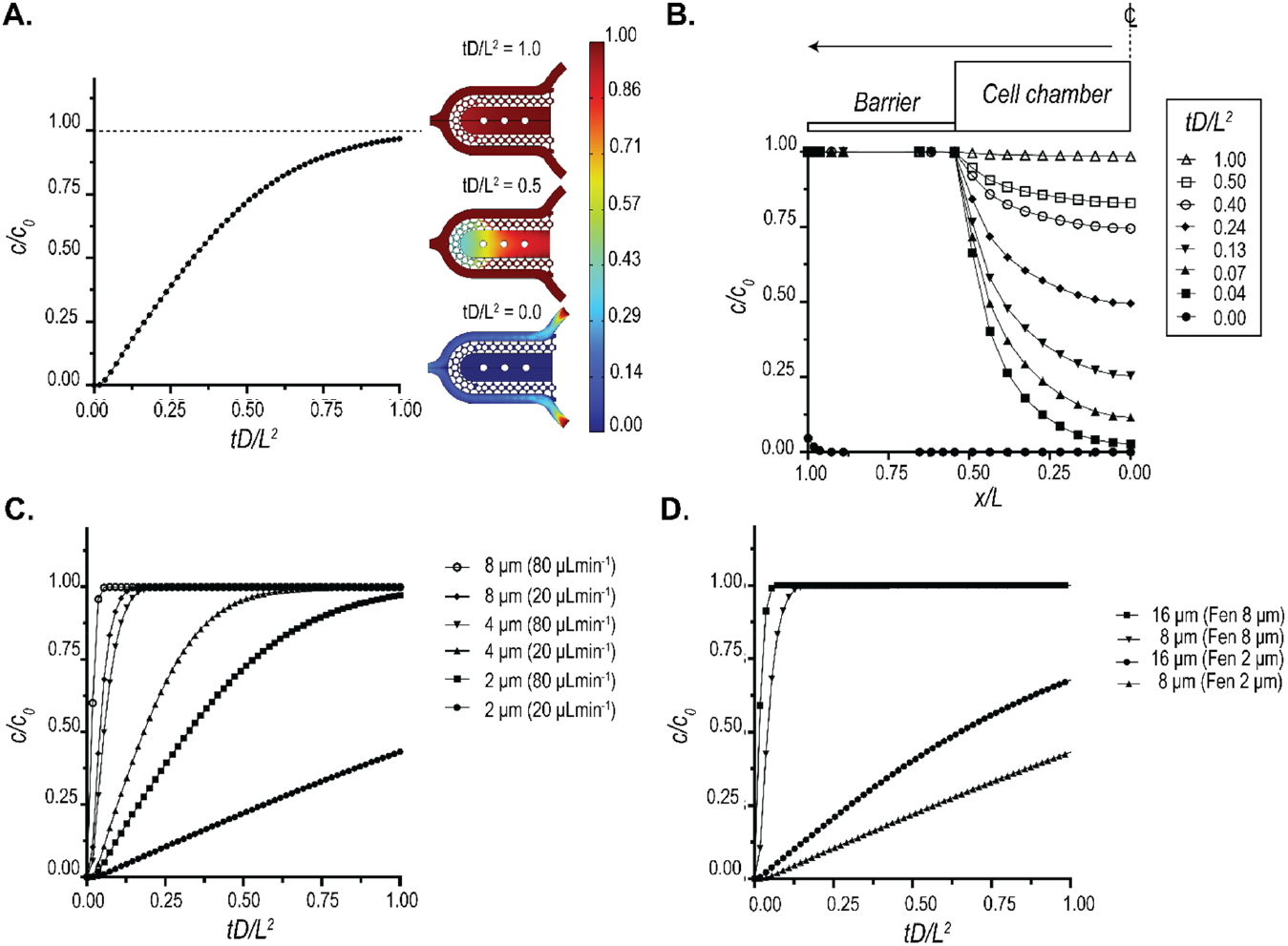
Characterization of biomolecular diffusion across the pillar array barrier. **(A)** FEM prediction of the space-averaged concentration at the center of the chamber when a 7 mmol/m^3^ (c_0_) dose of 70 kDa dextran was injected into the media channel at a flow rate of 80 µL/min. The spatial distributions of dextran at representative time points are also shown. Non-dimensional values were plotted by using the diffusivity of 70 kDa dextran (D= 45.8 µm^2^/s) and characteristic length (L=275 µm). **(B)** Spatio-temporal evolution of 70 kDa dextran within the cell chamber for the same conditions as A. **(C)** Tuneability of diffusion offered by the pillar array as a function of fenestration height. The traces represent the space-averaged values of concentration at the cell chamber center. **(D)** Tuneability of diffusion offered by the pillar array as a function of the pore size. The traces represent the space-averaged values of concentration at the cell chamber center.

The value of 80 µL/min was used as the maximal rate of flow in our device used in the case of metabolic interrogation. Using this flow rate, we plotted the concentration profile of 70 kDa dextran within a section of the cell chamber at different time points (**Figure 3B**). The temporal concentration profiles reveal that even at such high flow rates, the transport of dextran is near-Fickian, suggestive of a primarily diffusion-based transport. Thus, our pillar array barrier protects the tissue from the shear forces of the perfusion medium and allows nutrient exchange via a diffusion-dominant transport, modeling an endothelial barrier in vivo.

The tunability of molecular transport can allow for modeling different disease states or accommodate the differential diffusion dynamics in different tissues.^24–27^ We demonstrate that this tunability can be very easily achieved using the pillar array barrier. **Figure 3C** shows the tunability of 70 kDa dextran diffusion into the cell chamber via modulation of the pillar height and flowrate. We observe a non-linear reduction in diffusion times when pillar height is increased. A similar change in diffusion times can be achieved by altering pore size and pillar height while keeping the flow rate constant, as shown in **Figure 3D** for a flow rate of 20 µL/min. These results demonstrate that the pillar array allows high modulation of molecular diffusion across the barrier, and can be designed such that in certain configurations, the transport across the barrier is diffusion dominant.

### Leveraging tunable diffusion barrier to design physiologically relevant cardiac microtissues

We used the pressure barrier functionality of the pillar array to aggregate hiPSC-derived cardiomyocytes in a cardiac MPS. The cell chamber has a width of 300 µm, while the media channels are 100 µm. The cell chamber and media channel heights are approximately 150 µm. The cell chamber consisted of 3 pillars (each 75 µm in diameter), separated by 225 µm (**Figure 4A**). Singularized hiPSC-derived cardiomyocytes were mixed with cardiac-specific hiPSC-derived fibroblasts in a ratio of 80:20 to form a cardiac microtissue. Details of the differentiation and loading process are provided in Methodology. Briefly, cell mixtures of cardiomyocytes and fibroblasts were loaded into the cell chamber via centrifugation at 300 r.c.f. for 3 minutes. The aggregation of cells offered by the combination of cell chamber dimension, the pillars in the cell chamber, and the barrier function afforded via the pillar array leads to densely condensed cardiac microtissue in the MPS (**Figure 4A**). The microtissue is 3D as validated by nuclei staining and high-resolution confocal imaging (**Figure 4A**). We further quantified the number of cells and the volume of microtissue over an extended time (∼months) and multiple biological replicates (**Figure 4B**). The mean values for the number of cells and tissue volume were 3685 (coefficient of variation/cv: 0.40) and 1.91×10^7^ µm^3^ (cv: 0.33), respectively. The cell density was calculated for each microtissue using the cell number and volume and had an average value of 1.95×10^14^ cell/m^3^ (cv: 0.26; **Figure 4B**), which matches the values observed in human heart muscle.^28^

**Figure 4:**
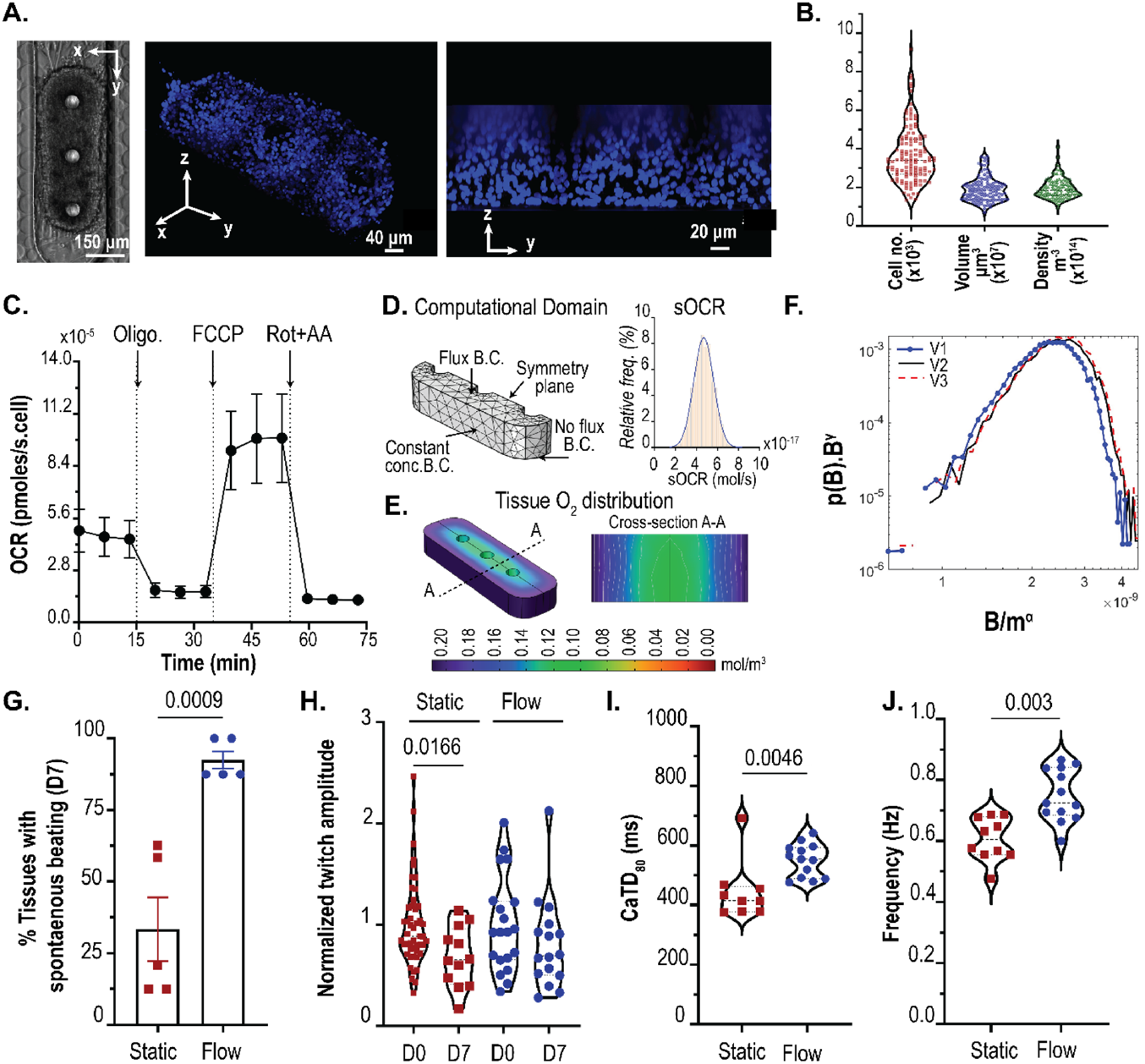
Pillar array allows the generation physiologically relevant microtissue. **(A)** 3D cardiac tissue formed within the MPS after aggregation of the single-cell. Shown are the brightfield image, and confocal images of the microtissue stained with nuclear stain DRAQ5. **(B)** Quantification of the cell number, volume, and density within the MPS. **(C)** Trace of mitochondrial oxygen consumption rates of cardiomyocytes in 2D obtained via Seahorse respirometer (n= 10; error bars represent SEM). **(D)** Computational domain used for the FEM PoM to predict oxygen profiles within the 3D tissue, and distribution of sOCR utilized in the reaction term. **(E)** Representative image of 3D oxygen profile obtained using the PoM. Shown here is the geometry V2. **(F)** Log-log distribution of collapse of the tissue OCR to determine the relationship between tissue OCR B and mass of microtissue m. γ = 0.125 and α = 0.8825-0.95, as determined by finite size scaling. **(G)** Comparison of percentage tissues beating after day7 in static vs. flow conditions. **(H)** Twitch amplitude comparison between day 0 vs day 7. Significance between day 0 and day 7 tested using student’s t-test. Comparison of calcium transient duration CaTD_80_ **(I)** and beating frequency **(J)** of microtissues on day 7 for static vs flow conditions.

After condensation, the microtissues in our MPS had mean dimensions of 243.8 µm x 621.6 µm x 124.7 µm (cv: 0.05, 0.28, and 0.95 respectively). We wanted to predict whether these dimensions of the microtissue would lead to the formation of necrotic cores, given the high cell density, and whether the microtissue remains metabolically active and viable. Thus, we combined oxygen consumption measurements obtained from 2D culture with a reaction-diffusion finite element computational model.

Briefly, we conducted oxygen consumption measurements across different batches of differentiation using an Agilent Seahorse XFe96 machine. Traces of oxygen consumption rate (**OCR**) were obtained at basal state and upon exposure to mitochondrial ATP synthase inhibitor oligomycin, protonophore FCCP, and rotenone and antimycin (Rot+AA) that inhibit complex I and III activities, respectively (**Figure 4C**).^29^ We measured a mean value of single-cell OCR (**sOCR**) of 4.67×10^−5^ pmoles/s upon normalization with the number of cells per well. At the same time, we see a huge variation in this sOCR (cv: 0.69). Our mean and variation of the sOCR matches with data reported elsewhere in the literature on hiPSC-derived cardiomyocytes.^30^ To account for this wide variation of sOCR, we implemented a population of model (PoM) approach to the reaction-diffusion model (detailed in Methodology). In this PoM, the reaction-diffusion model was solved 10,000 times, each time with a different value of sOCR that represented experimentally observed values. The computational domain (**Figure 4D**) representing the tissue was based on measurements of the tissue geometry.

We chose a base geometry (**V1**) whose dimensions were 243.8 µm x 621.6 µm x 62.35 µm, which are within the ranges of dimensions observed for our microtissue. Next, we doubled the height of V1 to obtain another geometry **V2** (243.8 µm x 621.6 µm x 124.7 µm). Since a higher variation was observed in the length of the tissue, we used another geometry (**V3**) of (288 µm x 950 µm x 124.7 µm). Overall, these geometries encompass the range of tissue volumes observed experimentally (V1: 0.8×10^7^ µm^3^; V2: 1.6×10^7^ µm^3^; V3: 3.1×10^7^ µm^3^). The boundary condition (**B.C.)** imposed on the top surface of the tissue was a flux term, capturing the diffusion of oxygen via the PDMS.^31^ A no-flux B.C. was imposed on the lower surface of the tissue facing glass. The side walls of the tissue were imposed with a constant oxygen concentration B.C. of 0.21 mol/m^3^. The distribution of sOCR, used in the reaction term R (see Methodology), to simulate the statistical nature of the tissue OCR is shown in **Figure 4D**. The PoM produced profiles of oxygen within the tissue and the tissue OCR (**B**), defined as the surface integral of the oxygen flux into the tissue. **Figure 4E** shows the oxygen concentration distribution in tissue geometry V3 with sOCR prescribed at 6.1×10^−17^ mol/s. Even with the higher end of the sOCR used, our PoM reveals that within the parameters used, the oxygen concentration profiles remain well above the critical value of 0.04 mol/m^3^ at which cells are believed to undergo cell death.^32^ **Figure S2A** shows the tissue OCR (or B distributions) for V1, V2, and V3. As expected, tissue OCR increased with increasing volume of the tissue. To investigate how these B distributions scale with size, we assumed the probability density function suggested by Zaoli et al.^33^ and shown in **Eqn. 1** as follows:

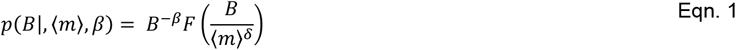

where, δ is the scaling exponent defining the relation between volume (or masses; m) and tissue OCR (B), and F is the general scaling function. The exponent β is a normalization exponent and is typically set to one.^34^ To see how these tissue OCRs scale with respect to mass, we performed a finite-size scaling collapse of the distributions.^35,36^ To measure the statistical distance between probability distributions, we leveraged the concept of probability contiguity as suggested by Bhattacharjee and Seno,^35,36^ and determined the scaling factor δ via a differential evolution optimization^37^ the leads to the best collapse of the B distributions for V1, V2, and V3. The collapse reveals a non-isometric scaling of tissue OCR (α = 0.8825) with respect to mass. These non-isometric scaling is indicative of near-diffusion limited tissue that are supposed to closely mimic physiological metabolism.^38–40^

Based on the OCR profiles predicted by our model, the microtissue should not undergo functional decay if there is sufficient diffusion of nutrients into the tissue. To test this, we compared our microtissues cultured in static vs under dynamic medium perfusion at 20 µL/h. This value of dynamic perfusion rate was chosen so that the shear stress at the walls was ∼ 1 dynes/cm^2^, which is considered near physiological.^41,42^ The hiPSC-derived cardiac tissue exhibit automaticity (i.e. spontaneous beating without electrical stimulation).^43^ To test the functionality of the microtissue, we used three metrics: twitch amplitude, calcium transient duration at 80% repolarization percentage (CaTD_80_), and beating frequency. Twitch amplitude is associated with the contractile nature of the cardiac tissue, i.e. how much the tissue contracts during a spontaneous beat. The CaTD_80_ provides a proxy measurement of the membrane potential waveform of the cardiac tissue and the beating rate provides the rate of spontaneous beating.^43^ We used published methods to measure the twitch amplitude, CaTD_80_, and the equivalent beating rate of the microtissues.^3,18,44^ We monitored the spontaneous beating of the hiPSC-derived cardiac tissue over 1 week/7 days. Microtissues cultured in static condition were fed every 2 days with fresh medium. Across different batches of measurements, 2/3^rd^ of the microtissues cultured in static conditions failed to beat spontaneously, indicative of functional degradation (**Figure 4F**). In contrast, 92% of the microtissues in dynamic perfusion had spontaneous beating, indicative of the functional integrity over 7 days. These microtissues with spontaneous beating at day 7 were considered functionally active tissues. Within these functionally active tissues, static culture led to the peak twitch amplitude of the microtissue being slightly reduced on day 7 (**Figure 4G**), whereas there was no statistical difference between the microtissues on day 0 vs day 7 in MPS where there was dynamic perfusion. The microtissues cultured in flow had higher CaTD_80_ and beating rate compared to static cultured microtissues (**Figure 4 H-I**). Thus, the pillar array allows the diffusion of nutrients and metabolites to maintain the functionality of physiologically dense 3D cardiac tissue.

### Pillar array as an engineered interface for creating heterotypic multi-tissue models

The pillar array can be leveraged to create an interface between endothelial cells (**EC**s) and the perivascular/parenchymal cardiac tissue (**Figure 5A**). Such heterotypic models usually require scaling of the different tissues and/or providing physiologically meaningful fluidic shear stress.^45^ To create a design that allows the scaling of two compartments with the pillar array as an engineered interface, we incorporated a modification of the two-step photolithography process for a three-step photolithography fabrication process (**Figure 5B**). Briefly, we spun coat a silicon master wafer with a desired thickness of photoresist. This thickness is the target pillar height for the barrier. After exposing the photoresist to patterns incorporating the pore size, the layer was developed to incorporate the barrier design on the master wafer. Next, another layer of a photoresist was used to generate the second layer (i.e. media channel) microstructure via the use of an alignment marker. After developing this layer, the third layer (i.e. cell chamber) microstructure was generated. The SEM for the master mold using this recipe is shown in **Figure 5C**. In this study, we created a barrier height of approximately 10 µm, that will allow cells to migrate between the media channel and the cell chamber. The media channels were approximately 50 µm, and cell chamber was approximately 100 µm. The corresponding polymeric PDMS replica mold with a pillar height of approximately 10 µm obtained from the master mold is shown in **Figure 5D**. The multi-layered fabrication demonstrated here will allow us to scale tissues and cell chambers to create heterotypic cell models in the future. To demonstrate one such heterotypic model, we created cardiac tissue consisting of cardiomyocytes and fibroblasts with ECs surrounding the cell chamber.

**Figure 5:**
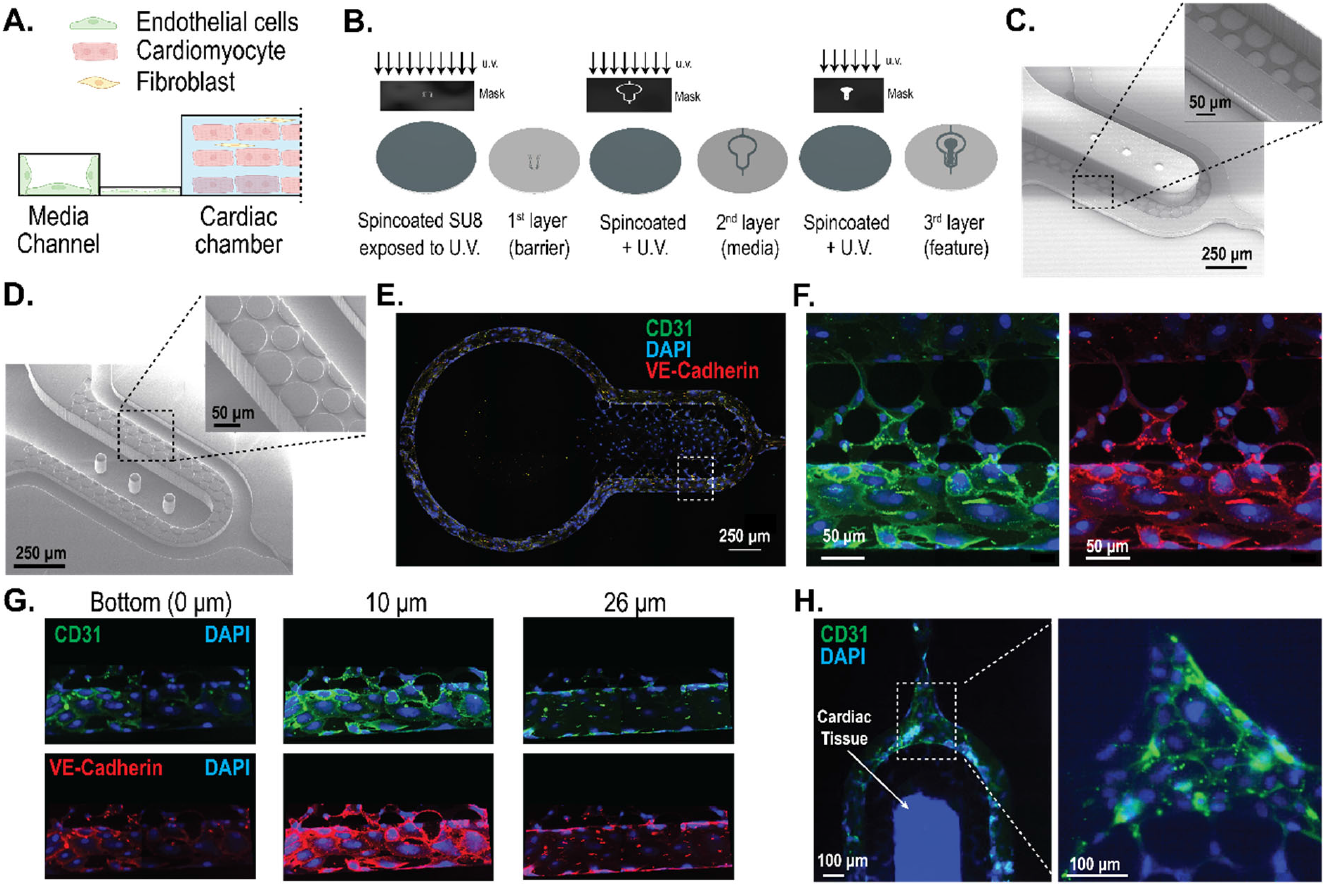
Engineering heterotypic cell models leveraging the interface created by the pillar array barrier. **(A)** A two-compartment model of endothelial cells and cardiac tissue, whereby the pillar array forms the engineered interface. **(B)** Fabrication schema for a two-compartment model with differential heights of the media channel and the cardiac chamber using a three-step/layer photolithography process to create master molds. **(C)** SEM image of the master mold with differential heights of SU8 microstructures for the two-tissue model. The media channel and cell chamber heights are 50 µm and 100 µm, respectively. The pillar height shown is 10 µm. **(D)** SEM image of PDMS replica mold obtained from the master mold. **(E)** Representative images showing endothelial cells forming a monolayer in the media channel cultured under near-physiological flowrates for 3 days. **(F)** High-resolution confocal image of endothelial cells from inset in (E), showing the ability to form cell layer across the pillar array barrier. **(G)** Confocal images of the vascular compartment at different heights show the coverage of the fenestration and media channel wall with endothelial cells. **(H)** Representative image of cardiac tissue, consisting of cardiomyocytes and fibroblasts, surrounded by endothelial cells in the media channel.

We cultured human coronary artery endothelial cells (**HCAEC**s) in the media channels of the cardiac MPS. Cells were loaded and cultured in static overnight after which fluid flow was introduced in the media channel. The flow rates were slowly ramped such that the shear stress experienced by the HCAECs for the first 12 hours were at ∼ 0.5 dynes/cm^2^, the next 24 hours at ∼ 1 dynes/cm^2^, and finally at ∼ 2 dynes/cm^2^ for the remaining 36 hours. Representative images of the HCAECs cultured under dynamic fluid flow conditions for 3 days in the cardiac MPS are shown in **Figure 5E**-**F**. We performed confocal imaging to assess how well the HCAECs formed a monolayer within the channel (as envisioned in **Figure 5A**) to add an active element to the passive barrier. We observed that the HCAECs covered the fenestrations, but did not form a 3D lumen covering all 4 walls of the channel (**Figure 5G**). We note that this is similar to any membrane-based barrier, where endothelial cells only cover the porous barrier.^8,46^ It is also noted that our pillar array diffusion barrier allows the formation of contiguous cell-cell interaction at the interface (**Figure 5F**), rather than discrete nature of contact using microchannel-based fenestrations.^14^ We demonstrate that the MPS platform consisting of the pillar array barrier can be leveraged to create heterotypic cardiac models. Briefly, we introduced cardiomyocytes and fibroblasts in the cell chamber and allowed aggregation for a day. The following day, we functionalized the media channels with fibronectin, followed by the introduction of HCAECs. **Figure 5H** shows the heterotypic cardiovascular model formed within the MPS after 3 days in culture. In vitro cardiac MPS consisting of heterotypic cellular components such as endothelial cells and macrophages have been proposed to create more physiologically relevant organ models and remains the scope of future investigation leveraging our MPS platform with tunable barrier interface.

## CONCLUSION

We report using standard multi-layered photolithography to create MPS devices with polymeric pillar arrays as tunable diffusion barriers. This microfluidic engineered interface acts both as a burst capillary valve and diffusion barrier to the tissue. We demonstrated the tunability of the barrier to control biomolecular diffusion via the alteration of the geometric parameters of the array. The cardiac tissue generated within these devices achieves physiologically dense tissue and maintains functionality afforded by the nutrient diffusion across the barrier at physiologically relevant dynamic flow rates. The pillar array can also be used to create tissue barrier models, such as vascularized cardiac tissue. The ease of fabricating multi-layered tunable diffusion barriers makes these pillar arrays attractive for creating scaling models for multi-tissue MPS platforms and creating sophisticated models for drug/therapy discovery and drug testing.

## ACKNOWLEDGEMENTS

This research was supported by NIH UH3DK120004 and the NSF Engineering Research Center for Advanced Technologies for Preservation of Biological Systems (ATP-Bio) NSF EEC 1941543.

## AUTHOR CONTRIBUTIONS

I.G. and K.E.H. conceptualized the study. I.G. conducted the computational modeling and analysis. I.G. and Y.K. conducted microfabrication and characterization. G.N., B.S., and T.N. optimized the cardiac tissue composition within the MPS and the imaging. J.V. conducted the Seahorse experiments. I.G., Y.K., and K.Y. optimized endothelial cell loading in the media channels. I.G., Y.K., G.N., B.S., J.V., K.Y., T.N., K.E.H. analyzed the data. I.G. drafted the manuscript, and all authors edited it.

## COMPETING INTERESTS

K.E.H. and B.S. have a financial relationship with Organos Inc. The remaining authors declare no competing interests.

## METHODOLOGY

### Fabrication

The cardiac MPS was fabricated using photolithography and casting of polydimethylsiloxane (**PDMS**). Briefly, the design of the MPS device was made using a computer-aided design software package (AutoCAD, Autodesk Inc., San Rafel, CA). The design consisted of two layers: the fenestration layer and a feature layer consisting of the cell chamber with media channels. These designs were used to create two 5”x5” laser plotted photomasks, which were emulsions printed onto a transparency, with a resolution of 50,800 DPI (EMS Thin Metal Parts, Colorado Springs, CO). For photolithography, a 100 mm diameter Si wafer was first cleaned using piranha solution (1:3 v/v mix of H_2_SO_4_:H_2_O_2_), followed by spin-coating of a thin layer of negative photoresist (SU8 2002/2005, Kayaku Advanced Material, Westborough, MA) whose height was equal to that of the desired height of the fenestration layer. After soft baking, the photoresist was exposed to UV light using the fenestration photomask followed by a post-exposure bake. The resist was then developed in propylene glycol methyl ether acetate followed by a hard bake at 180*°C*. A next round of spin-coating of a thicker negative photoresist (SU8 2100, Kayaku Advanced Material, Westborough, MA) was then performed for a desired height of the cell chamber. After soft baking, the Si wafer was aligned to the feature layer photomask using fiduciary markers to align the fenestration and cell chamber layers. The photoresist was exposed to UV light and then developed after a post-exposure bake. The process of hard baking was performed, followed by passivation of the Si mold by exposing it to trichloro [1H, 1H, 2H, 2H-perfluorooctyl] silane (Sigma-Aldrich, catalog 448931) overnight via vapor deposition. This master mold was used to obtain replica molds of PDMS by pouring 20 g of de-gassed 10:1 mixture of PDMS oligomer and cross-linking agent (Sylgard 184, Dow Corning, Midland, MI) and curing for 8 hours at 65*°C* in an oven. PDMS stamps were peeled off after cooling to room temperature, and 0.75 mm holes punched at the loading and media port positions using biopsy punch (Ted Pella, Redding, CA). PDMS stamps were then exposed to oxygen plasma (Plasma Equipment Technical Services, Livermore, CA) for 60s (power: 21 W; flow: 98.8 sccm; pressure: 20 mTorr) and bonded to glass to create the device.

### SEM characterization

Gold/palladium was deposited on the surface of PDMS samples for 100 s using the sputter coater (Cressington, UK). FEI Quanta 3D FEG SEM (Field Electron and Ion, Hillsboro, OR) was used to acquire images.

### Finite element model

We used COMSOL 6.2 (COMSOL, Inc., Burlington, MA) to create finite element models (**FEM**s) of our cardiac MPS. We used FEM to quantify burst pressure in the diffusion barrier, the biomolecular transport, and the metabolism of the cardiac tissue within the MPS. Details of the FEM for each of the characterizations are provided in the following.

### Burst pressure calculations

We implemented a level set method available via COMSOL FEM package to simulate the air and water multiphase transport and calculate the burst pressures across different barrier designs. Geometries used for the simulation are shown in **Figures 2A & 2E**. We implemented a pressure boundary condition at the inlet to simulate the required pressure field to burst the capillary valve formed by the diffusion barrier. Outlet gauge pressure was set to 0 Pa. The properties for air and water were imported from the built-in material library within COMSOL. Details of the implementation within the software can be found in the documentation from COMSOL Inc. A laminar flow solver for Navier-Stokes equations incorporating surface tension forces was used to estimate the transport of mass and momentum for the two phases. Slip boundary condition was implemented by providing the contact angle between the material consisting of the walls (e.g. PDMS) and water using the Wetted Wall coupling feature in the software. Note that the contact angle of the solution domain boundaries can be determined experimentally by measuring the advancing contact angle in a goniometer. The multiphysics coupling between the level set and laminar flow solvers for the calculation of the water-air interface was performed via a segregated solver that involved a sequential solution of the two solvers.

### FEM characterization of biomolecular diffusion across barrier

Transient flow profiles of candidate biomolecule (dextran) within the MPS were estimated through the solution of incompressible Navier-Stokes equation utilizing the Laminar Flow module. Similarly, transient concentration profiles were estimated via the deployment of Transport of Diluted Species module. The two module solvers were coupled via the Multiphysics solver Reacting flow, Diluted species. For the fluid flow solver, the boundary conditions were set as mass flow rate and outlet was set as pressure outlet boundaries, while no-slip condition was set for rest of the boundaries. For the calculation of concentration profiles, inlets were prescribed with a concentration of the candidate biomolecule, and outlets were set as outflow boundary conditions, while no-flux condition was set for the rest of the boundaries.

### In Silico generation of cardiac tissue metabolic profiles in the MPS

We implemented reaction-diffusion FEM to predict the oxygen profile within the cardiac microtissue. Specifically, the reaction-diffusion equation was of the form:

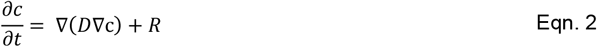

In this Eqn. 1, c is the concentration of oxygen, D the diffusion of oxygen in tissue, *∇* the standard del operator 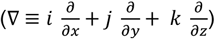, and R is the reaction rate that captured the oxygen consumption. In our study, we modeled R based on a Michaelis-Menten type schema, i.e.

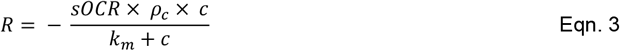

The values for single-cell oxygen consumption rate (**sOCR**) and cell density (***p***_*c*_) were obtained from experimental measurements (Results and Discussion). The value of the Michaelis-Menten constant (**k**_**m**_) was set at 6.9×10^−3^ mol/m^3^, based on value used for cardiac tissue elsewhere in the literature.^47–50^ Upon solving for steady-state profiles of oxygen in the microtissue, the OCR of the tissue was calculated as surfce integral of the inward flux of oxygen at the boundaries. To implement the inherent variability in cell metabolism arising from intraspecific differences or batch-to-batch variation of stem cell differentiation to cardiomyocytes, we implemented a population of model (**PoM**) approach whereby we imposed variation in sOCR based on experimentally observed values.

### Cardiac differentiation, culture, and cell loading into MPS

Cardiomyocyte cells were derived from human induced pluripotent stem cells (hiPSC). The hiPSC line WTC-11 was expanded on growth factor-reduced Matrigel-coated plates (Corning, 354248) in mTeSR1 Plus medium (Stemcell Technologies, 100-0276) that was changed daily, passaged at 80% confluency using Accutase (Thermo Fisher Scientific, A1110501), and plated at a density of 12,000 cells/cm2. Cells were fed culture medium supplemented with 5 μM Y-27632 dihydrochloride (Biogems, 1293823) for the first 24 h after passaging. Once confluent, the hiPSC cells were differentiated into human iCMs utilizing a chemically defined cardiomyocyte differentiation protocol with some modifications (https://doi.org:10.1073/pnas.1200250109). Briefly, hiPSCs were treated with 6 μM CHIR99021 (Biogems, 2520691) for 2 days in RPMI 1640 (Thermo Fisher Scientific, 11875119) with B27-insulin (Thermo Fisher Scientific, A1895601). The cells were subsequently treated with Wnt inhibitor IWP4 (Biogems, 6861787) in RPMI/B27-for another 2 days. Between 5–11 days of differentiation, RPMI/B27-medium was used and changed every other day. A robust spontaneous contractile activity was typically observed on days 8-10 of differentiation, at which the medium was switched to RPMI/B27+insulin (Thermo Fisher Scientific, 17504044). Cardiomyocyte purity was characterized using flow cytometry for cardiac troponin T (cTnT).

Once spontaneous contractile activity was observed, the hiPSC-CMs were dissociated with TrypLE 10x (Thermo Fisher Scientific, A1217703) and suspended in EB20 medium supplemented with 10 µM Y27632. To create isogenic iCM-icFb micromuscles in the MPS, we purified the iCMs by replating at a density of 100,000 cells/cm2 onto Matrigel, culturing in RPMI/B27+ without glucose (Thermo Fisher Scientific, 11879020) with 5 mM sodium Lactate (Sigma Aldrich, 71718) for 4 days. Cells were allowed to recover in RPMI/B27+ for 2 days. In parallel, we generated hiPSC-derived cardiac fibroblasts following a previously published protocol (https://doi.org:10.1007/7651_2020_300). On days 12-14, iCMs were dissociated using TrypLE10X, and icFbs were dissociated using Accutase. An isogenic cardiac microtissue was created by mixing 80% iCM-20% icFb EB20 by suspending cells with a density of approximately 2×10^6^ cells/mL. 15,000 cells/8 μL that was injected into the loading port of each MPS. After 3 min of centrifugation at 300g, MPS were inspected under the microscope. Chambers that were not filled with iCMs at this point were discarded. MPS were fed with 200 μL EB20 medium supplemented with 10 µM Y27632 into the inlet tip, and gravity allowed for constant flow to the outlet until equilibrium was reached. The following day and every day from then on, the medium was changed to 80% RPMI/B27+, 20% FGM3.

### Image Acquisition for Calcium Transient and Contractile Activity Studies

Calcium transient and beating physiology studies were conducted using high-resolution image acquisition techniques. Cardiac MPS (microphysiological systems) were maintained at 37°C during imaging, using a Tokai Hit stage with integrated heating. Spontaneous recordings included 6-second fluorescent videos (using the GCaMP6f WTC hiPSC line) for calcium transient analysis and 6-second brightfield videos to evaluate contractile activity. Post-experiment analysis was performed using a custom Python library, developed in-house, capable of processing fluorescence intensity over time and quantifying contractile motion from the brightfield recordings. Imaging was carried out with a NIKON TE300HEM microscope paired with a HAMAMATSU C11440/ORCA-Flash 4.0 digital CMOS camera, capturing videos at 100 frames per second (FPS). For fluorescence imaging, the Lumencor SpectraX Light Engine (Beaverton, OR) was employed in combination with a QUAD filter from Semrock (IDEX, Rochester, NY). Video acquisition was managed using Nikon’s NIS-Elements software.

### Endothelial cells culture and cell loading into MPS

Human coronary artery endothelial cells (PromoCell, Heidelberg, Germany) were cultured in the endothelial cells growth medium MV containing the supplements kit (PromoCell Heidelberg, Germany), maintained at 37 C in a 5 % CO2 incubator. The MPS was functionalized with 0.5 mg/mL of fibronectin (Sigma, Burlington, MA) for 1 hour at 37 C before loading cells. A concentration of 30 ×10^6^ cells/mL was loaded into the media ports of MPS using a pipette. After 1-2 hours, non-attached cells were washed out by flushing medium. For culturing within the MPS, medium used was 80% RPMI/B27+, 20% FGM3 supplemented with 1 ng/mL VEGF.

### Immunofluorescent imaging

MPS were flushed with PBS via the media channel for 10 min, after which tissues were fixed with 4% paraformaldehyde by 15 min exposure followed by PBS wash (2x). For staining the cardiac tissue, the devices were cut clean using a scalpel to expose the tissue, which is still attached to the PDMS. Following this tissues were stained by submerging the PDMS and tissue in different staining solutions. Tissues were first blocked with blocking buffer (1% BSA, 10% FBS, 0.5% Triton, 0.05% sodium azide) overnight at 4 °C. The next day, they were submerged in primary antibodies (mouse anti α-actinin, Life technologies 41811; rabbit anti-myosin light chain 2V (MLC-2V), Proteintech 10906-1-AP) at 1:100 concentration in blocking buffer for 48 h at 4 °C. Tissues were then washed twice at 25 °C in blocking buffer for 2–3 h and washed a third time at 4 °C overnight. The secondary antibodies (goat anti-mouse IgG Alexa 568 H+L, Life Technology a11004; goat anti-rabbit IgG Alexa 488 H+L, Life Technology a11008) along with 1:600 DRAQ5 (Abcam, ab108410) were incubated in blocking buffer for 24 h. Tissues were then washed twice at 25 °C in blocking buffer for 2–3 h and a third time at 4 °C overnight before tissues were imaged.

### Mitochondrial respiration measurements

Oxygen consumption of hiPSC-derived cardiomyocytes were performed in a Seahorse XFe96 machine (Agilent Technologies, Inc., Santa Clara, CA) using previously reported protocol.^29^ Briefly, approximately 20,000 cells were seeded in each well and incubated overnight in a cell culture incubator at 37°C and 5% CO_2_. After overnight incubation, samples were incubated with the assay buffer (Seahorse XF base medium, 1 mM pyruvate, 10 mM glucose, and 2 mM glutamax) in a non-CO2 incubator for 45 minutes. After which they were washed again and mitochondrial respiration via oxygen consumption rate (OCR) was measured in the XFe96 plate reader. Measurements were made at basal state for 15 mins. Injections of 1.5 *μ*M ATPase inhibitor oligomycin, 2 *μ*M protonophore FCCP, and 0.5 *μ*M mixture of ETC complex III inhibitor antimycin-A and ETC complex I inhibitor Rotenone were made to measure the mitochondrial response. All concentrations reported are the final concentrations in each well.

### Statistical analyses

The software GraphPad Prism (GraphPad Software, San Diego USA) was used for statistical analyses. The statistical differences between multiple groups were compared using one-way analysis of variance (ANOVA) followed by post hoc Tukey HSD to find means that were significantly different from each other. Differences between means of two sample data were tested by Student t-test.

## SUPPLEMENTARY

**Figure S1:**
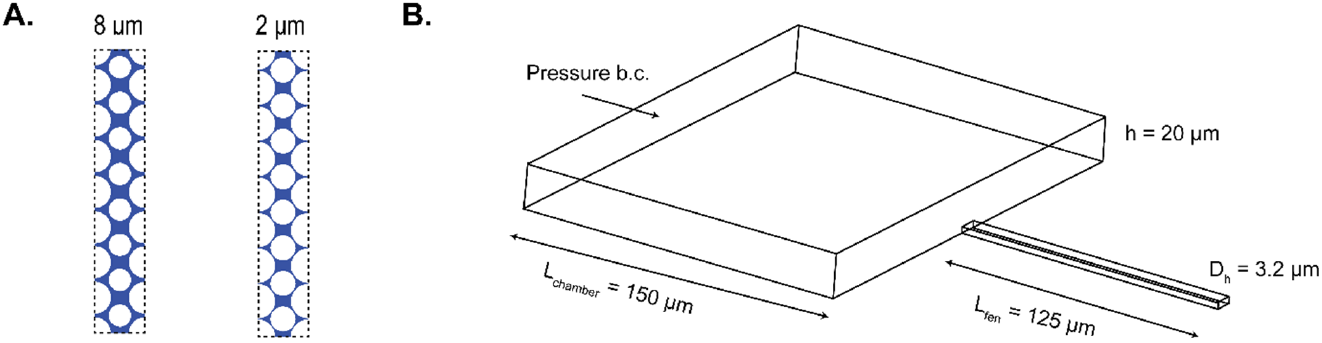
**(A)** Variation of pore size as defined by the distance between the pillars changes the porosity of the pillar-based fenestration layer. Shown here are two interfaces created in a 125 µm x 708 µm rectangle with 8 µm and 2 µm pore sizes and porosity of 31% and 19%, respectively. **(B)** A single microchannel-based fenestra finite element model was used to simulate the movement of the air-water interface across the microfluidic barrier to quantify burst pressure.

**Figure S2:**
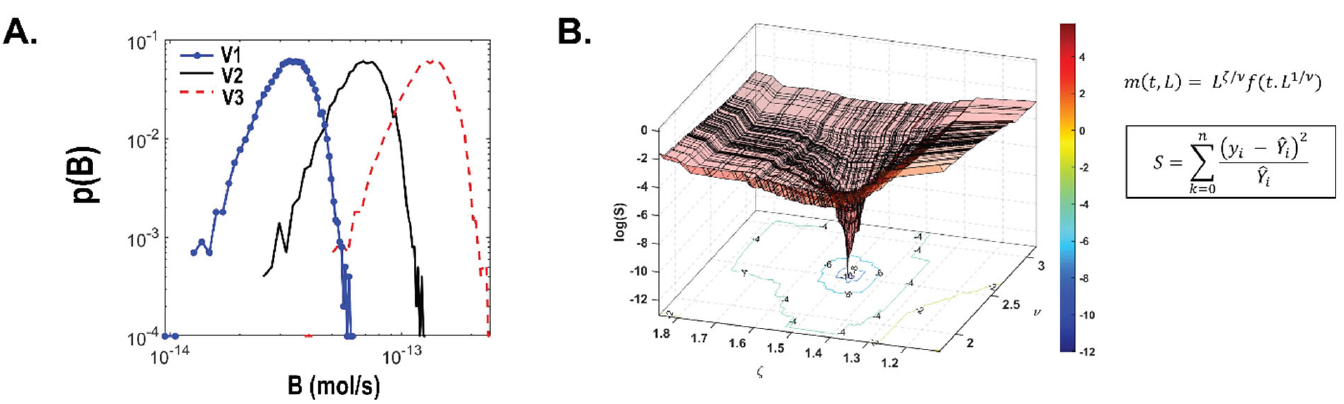
**(A)** Tissue OCR B probability distributions obtained for V1, V2, and V3 using FEM PoM method. **(B)** Differential evolution approach was used to obtain parameters for probability collapse. Show here is the estimation of parameters ν andζ for the collapse of probability distributions of m to estimate the scaling with respect to L and t, described by a scaling function f.

